# Miniscope Recording Calcium Signals at Hippocampus of Mice Navigating an Odor Plume

**DOI:** 10.1101/2024.06.12.598681

**Authors:** Fabio M. Simoes de Souza, Ryan Williamson, Connor McCullough, Alec Teel, Gregory Futia, Ming Ma, Aaron True, John. P. Crimaldi, Emily Gibson, Diego Restrepo

## Abstract

Mice navigate an odor plume with a complex spatiotemporal structure in the dark to find the source of odorants. This article describes a protocol to monitor behavior and record Ca^2+^ transients in dorsal CA1 stratum pyramidale neurons in hippocampus (dCA1) in mice navigating an odor plume in a 50 cm x 50 cm x 25 cm odor arena. An epifluorescence miniscope focused through a GRIN lens imaged Ca^2+^ transients in dCA1 neurons expressing the calcium sensor GCaMP6f in Thy1-GCaMP6f mice. The paper describes the behavioral protocol to train the mice to perform this odor plume navigation task in an automated odor arena. The methods include a step-by-step procedure for the surgery for GRIN lens implantation and baseplate placement for imaging GCaMP6f in CA1. The article provides information on real-time tracking of the mouse position to automate the start of the trials and delivery of a sugar water reward. In addition, the protocol includes information on using of an interface board to synchronize metadata describing the automation of the odor navigation task and frame times for the miniscope and a digital camera tracking mouse position. Moreover, the methods delineate the pipeline used to process GCaMP6f fluorescence movies by motion correction using NorMCorre followed by identification of regions of interest with EXTRACT. Finally, the paper describes an artificial neural network approach to decode spatial paths from CA1 neural ensemble activity to predict mouse navigation of the odor plume.

**SUMMARY:** This protocol describes how to investigate the brain-behavior relationship in hippocampal CA1 in mice navigating an odor plume. This article provides a step-by-step protocol, including the surgery to access imaging of the hippocampus, behavioral training, miniscope GCaMP6f recording and processing of the brain and behavioral data to decode the mouse position from ROI neural activity.

## INTRODUCTION

Although significant progress has been made understanding neural circuits involved in olfactory navigation in head-fixed mice^1–3^, and navigation strategies in freely moving mice^4–8^ the role of neural circuits in ethologically relevant freely moving navigation of turbulent odor plumes is still unknown. This article describes how to monitor neural activity by imaging Ca^2+^ transients in cells expressing the genetically encoded calcium sensor GCaMP6f in Thy1-GCaMP6f mice^9^. to study whether sequential neural dynamics of dorsal CA1 stratum pyramidale neurons in the hippocampus (dCA1) plays a role in odorant plume navigation. The methods provide information on imaged GCaMP6f fluorescence through a miniature epifluorescence microscope focused through a GRIN lens on dCA1^10–12^. The methods explain how to monitor simultaneously spatial navigation and dCA1 neuron GCaMP6f calcium transients in mice performing an odor-plume navigation task where they received a water reward when they reached the spout delivering an odorant into an odor arena with a background laminar air flow^13, 14^. This article describes the methods required to achieve this task (Figure 1), including the stereotaxic surgery for the implantation of GRIN lenses, the placement of a baseplate to secure the miniscope to the skull in a freely moving mouse, imaging with the miniature microscope and monitoring mouse movement with a high-speed digital camera, data pre-processing for removing motion artifacts and finding the regions of interest (ROIs), and preparation of datasets and artificial neural network training and prediction for decoding the X and Y positions of the mouse in the odor arena from changes in fluorescence in dCA1 ROIs^7^.

**Figure 1.**
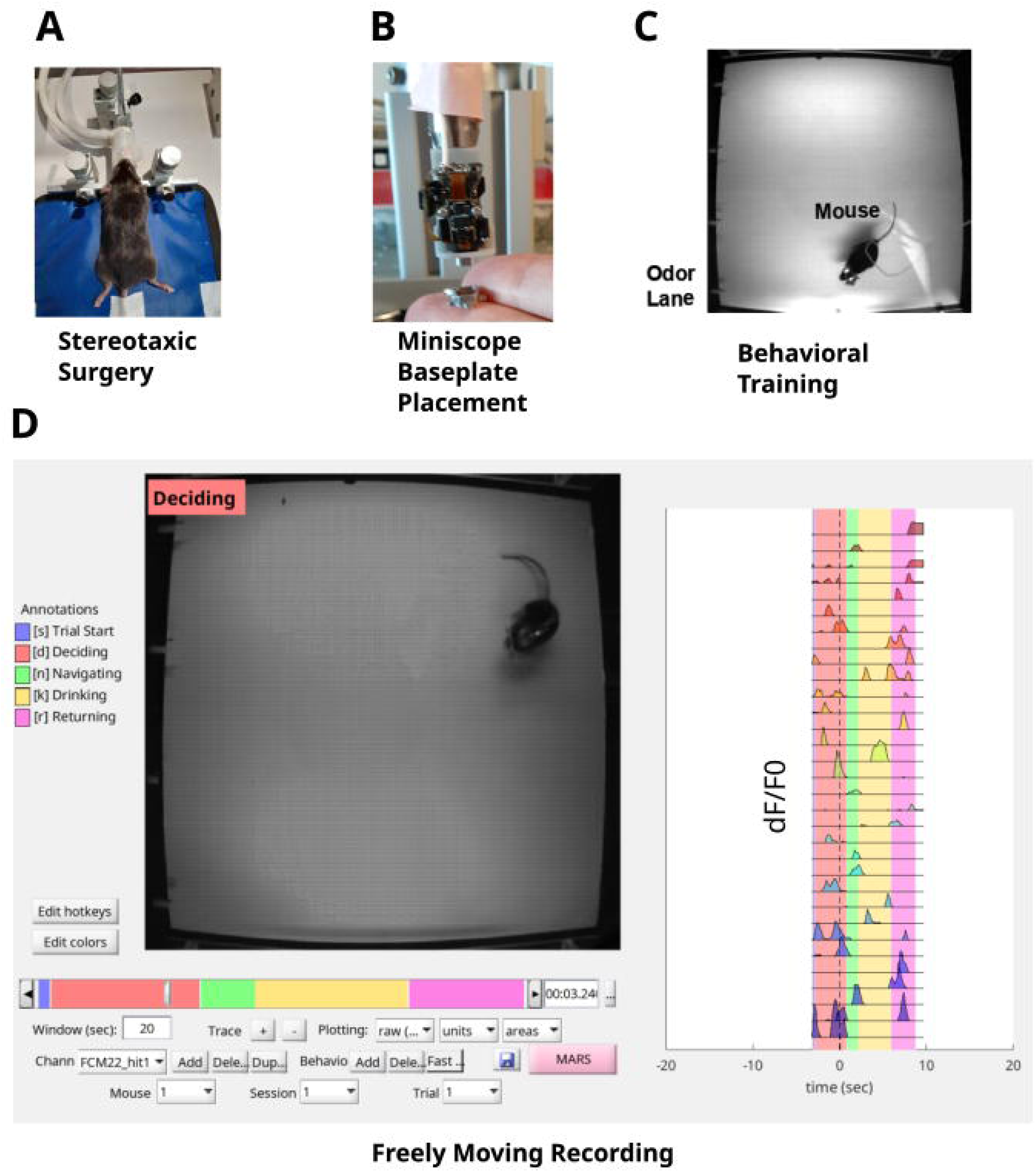
Study Pipeline. A: The stereotaxic surgery consists of implanting a GRIN lens in the CA1 layer of hippocampus and a headplate on the skull for head fixing the mouse. B: A baseplate is placed on the top of the GRIN lens to allow accessing optically the fluorescence of the CA1 neurons with a miniscope. C: The mouse is trained on the odor plume navigation task. D: Freely moving recording of the mouse behavior and CA1 neurons navigating the odor plume.

Miniscope recording of calcium signals in the CA1 region of the hippocampus of mice navigating an odor plume is relevant for understanding the computation of neural circuits involved with olfaction and spatial information in the complex behavior task of odor-plume navigation ^2, 14–16^. The CA1 region of the hippocampus plays a role in spatial navigation and is crucial for creating a cognitive map of the environment for efficient navigation ^17, 18^. Recording calcium signals with a miniscope is a valuable way to investigate the CA1 neurons that encode spatial information during odor plume navigation.

This technique combines the advantages of miniscope technology for recording GCaMP calcium signals with the well-established role of the CA1 hippocampus in spatial navigation to understand better how neural circuits drive complex behaviors ^19^. Alternatively, approaches using 2-photon microscopy can record CA1 neurons ^9, 20^, which requires a head-fixed mouse and restrains the possibility of freely moving to navigate an odor plume^21^. Local-field electrophysiological recordings of CA1 neurons allow the investigation of freely-moving mice navigating odor plumes^22^. Still, local field electrical signals impose limitations to estimating intracellular firing by isolating single-unit signals through spike sorting techniques. Miniscope signals allow the identification of ROIs associated directly with intracellular calcium signals in a reliable way ^10, 11^ to investigate neural computations at single-cell resolution precisely. Miniscope technology provides a unique opportunity to better understand how the CA1 region encodes spatial information based on odor cues.

Furthermore, this technique investigates how specific neuronal populations process odor information for navigation and the relationship between neuronal activity patterns and decision-making during odor plume tracking. This method can contribute to a better understanding of how the brain processes odor and spatial information. While miniscopes offer a single-cell resolution for recording a freely moving mouse’s brain, they require specialized surgery and data analysis expertise. In this paper, we provide a comprehensive protocol for helping researchers go through each step to investigate the neural mechanisms of odor-plume navigation.

The odor navigating task is a promising framework for studying neural coding and spatial odor cue memory in mice. The article’s findings indicate that it is possible to decode the trajectory of the mouse navigating an odor plume based on neuronal ensemble calcium signals in dCA1. Understanding the role of dCA1 calcium signals in odor plume navigation is a crucial step to crack the neural circuit basis for odor-guided navigation in realistic environments^13, 14^.

## PROTOCOL

Studies were carried in 3-to-6-month-old male and female Thy1-GCaMP6f transgenic mice^23^. All experimental protocols were approved by the Institutional Animal Care and Use Committee of the University of Colorado Anschutz Medical Campus, in accordance with National Institutes of Health guidelines. The surgical procedures for GRIN lens implantation (Protocol 1) and the baseplate placement (Protocol 2) were adapted from previous works^9, 24–29^.

1. Stereotaxic surgery for implanting a GRIN lens implantation in the hippocampus.

1.1 The surgeon wears sterile gloves, a headcover, surgical mask and a lab coat.

1.2 Sterile conditions are used during all survival surgeries. Sterilize all surgical equipment by autoclaving.

1.3 Place sterile drapes around the surgical area

1.4 Anesthetize the mouse by placing it in the induction chamber with 3% isoflurane for 10 min.

1.5 Check whether the mouse stops moving and is under deep anesthesia confirmed by pinching the hind paw to test the hind paw reflex.

1.6 Switch the isoflurane flow to the nose cone (Supplementary File 1).

1.7 Make sure the low-flow isoflurane anesthesia system for mice (Jove Materials Spreadsheet) has the temperature pad set to homeothermic settings and make sure the vapor source is set to external compressed.

1.8 Place the anesthetized mouse in the digital stereotaxic instrument (Jove Materials Spreadsheet). Place the front teeth of the mouse into the bite bar and place the nose cone in front of the nose to ensure the isoflurane flow and to lock the head into place. Adjust the level of anesthesia to between 2 to 3% according to the paw reflex after pinching (Figure 2 A).

**Figure 2:**
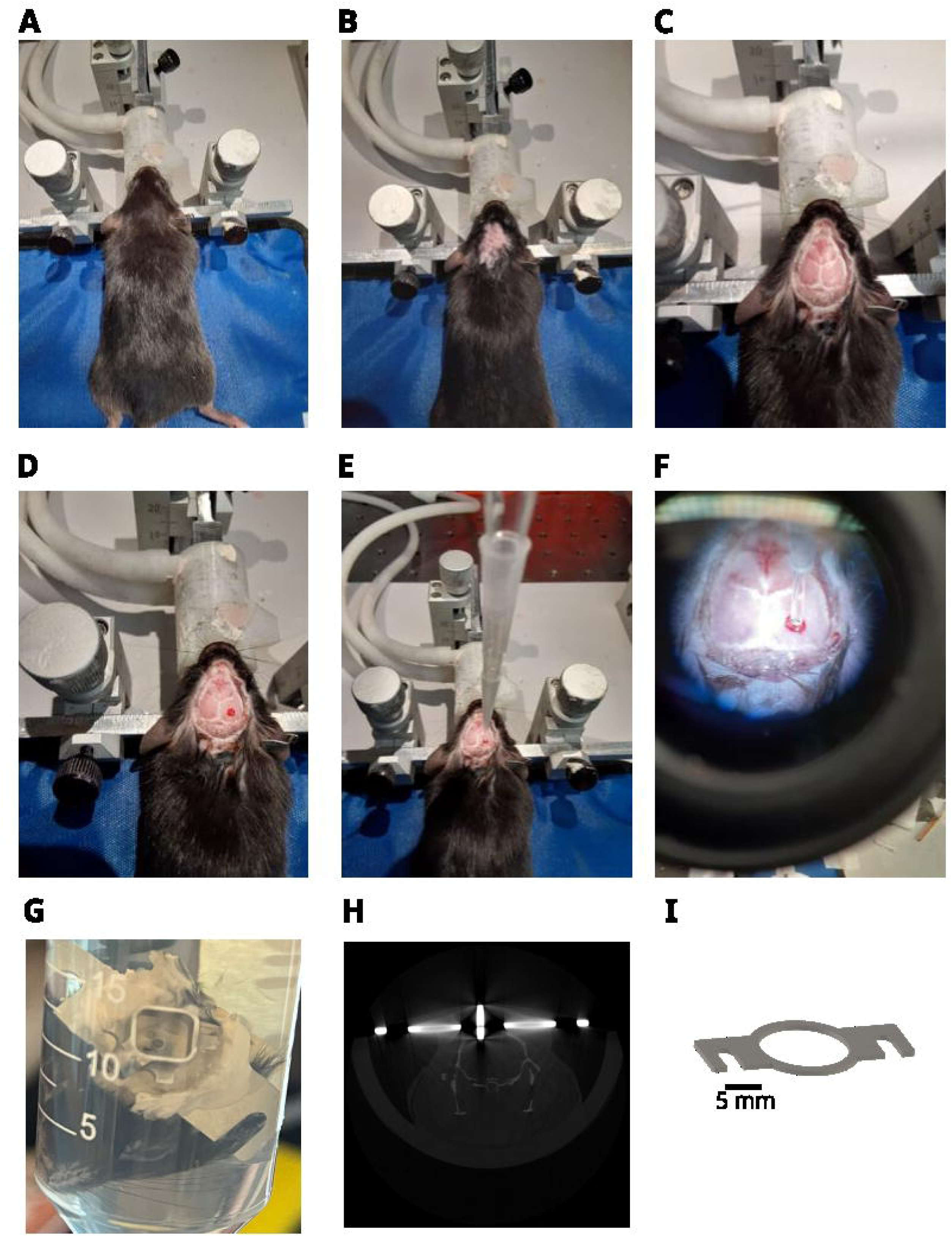
Stereotaxic Surgery. A: Place the anesthetized mouse in the stereotaxic apparatus. Adjust the level of anesthesia between 2 to 3% accordingly with the pinching paw reflex response B: Shave the hair above the head. C: Carefully drill a small permanent dent with drill to make a permanent dent on the top of the target location. D: Use the dental drill to open a circular perforation of 1.5 mm diameter to allow the implantation of a 1 mm diameter and 4 mm length GRIN lens into the Brain. E: Connect the GRIN lens holder on the micromanipulator and turn on the aspirator connected to the pipette to hold the GRIN lens. F: Implant the GRIN lens slowly into the cortex until reaches the depth of -1.25 mm below dura matter. G: Illustrative post-mortem fixed head showing the head-plate and base-plate cemented on the skull with the implanted GRIN lens. H: Post-mortem CT-scan of the head illustrating the Head Bar on the top of the cranium and the GRIN lens implanted inside of the cranium. I: Design of the head-plate to yield head-fixing the mouse.

1.9 Place the mouse ear bars into the ears and tighten. Ensure the mouse’s head is horizontal and does not move side-to-side or up and down. When the head is secure, lightly pressing downward on the skull shouldn’t cause the skull to slip out underneath the headbars.

1.10 Place the temperature probe in the rectum and tape the wire on the mouse’s tail to avoid movement. Set the temperature of the pad to maintain the rectal temperature at 37 degrees Celsius.

1.11 Add ophthalmic ointment to the eyes to prevent harmful air drying.

1.12 The hair is shaved when head is fixed and under the isoflurane flow provided by the nose cone to prevent movement. Shave the hair above the head with an electric shaver, then clear any remaining hairs off with hair remover and sterile q tip (Figure 2 B).

1.13 Make the operating field sterile by swabbing the scalp with a sterile Q tip with ethanol and betadine three times each.

1.14 Inject 0.1 mL of the local anesthetic lidocaine subcutaneously (S.Q.) under the skin between the eyes and ears with a 29 Gauge needle. This will make a bubble. Wait for a few minutes before cutting the skin with a scissors.

1.15 Using a tips only technique, pull the skin up carefully using forceps and use a pair of small scissors to remove a circular portion of the skin. Absorb any blood with a sterile cotton swab.

1.16 Once the bleeding stops, clean the skull with hydrogen peroxide (5%) using a sterile cotton swab. Bone landmarks, including where the sagittal suture and the bregmatic suture intersect to mark bregma should be easy to identify after this procedure.

1.17 Confirm that the skull is flat relative to the stereotaxic apparatus by making sure that bregma and lambda sutures are exactly at the same Z coordinate. This procedure is performed by placing a needle in the manipulator arm and verifying if the tip of a needle is at the same Z coordinate when touching both sutures. Otherwise, adjust the angle of the head by moving the bite bar in the Z plate and repeating the procedure until the skill is in a flat position.

1.18 Attach a pipette to the micromanipulator holding a needle to apply tattoo ink. Make a mark on the top of Bregma.

1.19 Zero the micromanipulator coordinates on Bregma and move to the coordinate above CA1 in the right hippocampus (medium-lateral +1.8 mm from Bregma, anterior-posterior -2.4 mm from Bregma)^2^. This is the location chosen by Radvansky and Dombeck to study vitural odor gradient navigation in mice^18, 30^. Carefully drill a small permanent dent of 1 mm diameter with a drill at speed of 10K min^-1^-(Osada, Jove Materials Spreadsheet) to make a permanent dent on the top of the target location (Figure 2 C).

1.20 Use the dental drill to open a circular perforation of 1.5 mm diameter to allow the implantation of a 1 mm diameter and 4 mm length GRIN lens into the Brain (Jove Materials Spreadsheet) (Figure 2 D).

1.21 Puncture the dura and cortex by inserting a 23 gauge needle in the middle of the hole and move the needle into the brain slowly (∼0.1 mm/min) until it reaches the depth of 1.25 mm below dura. Remove the needle slowly from the brain (∼0.1 mm/min). Use sterile cotton swabs and saline to clean the blood.

1.22 This protocol uses a custom made UCLA’s GRIN lens holder constructed with two 1ml micropipette tips cut to fit inside each other (GRIN lens holder, Jove Materials Spreadsheet) (Figure 2 E, F). Connect the GRIN lens holder on the micromanipulator and turn on the aspirator connected to the pipette to hold the GRIN lens. The air pressure produced by air suction holds the GRIN lens in place. This protocols uses a 4 mm length, 1 mm diameter GRIN lens. Implant the GRIN lens slowly into the cortex until reaches the depth of -1.25 mm below dura.

1.23 Place a drop of liquid tissue adhesive (Jove Materials Spreadsheet) into the circular cranium perforation to seal the hole. Wait a few minutes to dry.

1.24 Prepare quick adhesive cement -biocompatible methacrylate resin- (Jove Materials Spreadsheet) on the base of the GRIN lens to permanently seal it on the skull. Wait a few minutes and let it dry.

1.25 Turn off the aspirator to release and slowly remove the GRIN lens holder from the top of the GRIN lens that at this point is permanently attached to the skull.

1.26 Place the head bar on the top of the skull centering the hole on the GRIN lens (Figure 2 G, H, I) (Supplementary File 2).

1.27 Carefully place quick drying adhesive cement (Jove Materials Spreadsheet) around the base of the GRIN lens to seal the cranial window and attach the head bar to the skull. Place drying adhesive cement (Jove Materials Spreadsheet) in the middle of the head bar and on the sides, to keep it tightly fixed on the skull. Make sure it is completely dry. Ensure that no adhesive cement is placed on top of the GRIN lens otherwise it will permanently degrade the optical path and block the image.

1.28 Place a drop of low toxicity silicone adhesive (Jove Materials Spreadsheet) on top of the GRIN lens for protection from physical damage.

1.29 Turn off the Isoflurane

1.30 Inject buprenorphine SR subcutaneously 0.001 mg/g immediately after surgery for analgesia.

1.31 Take immediate post-operative care monitoring until the animal wakes from anesthesia. Post-operative care is taken until the fully recovery from the surgery by monitoring the mice for signs of pain. The animal that undergone surgery is not returned to the company of other animals. Briefly, movement around the cage, eating and drinking and normal reaction to handling is used to ensure that they are not under pain or stress. If the animals show signs of severe neurological or tissue damage, which was out of the ordinary for the procedure, they will be humanely euthanized immediately.

2. Baseplate placement for the miniscope

The procedures for head-fixing the mouse start two weeks after the animal fully recovers from the surgery. The procedure for imaging dCA1 start three weeks after the surgery after the animal full recovers and the GCaMP6f signal becomes strongly visible.

A baseplate is fixed on top of the GRIN lens for optical access of dCA1 GCaMP6f fluorescence through the GRIN lens using a miniscope. This protocol utilized the miniscope version 4 -V4 (Miniscope V4, Jove Materials Spreadsheet).

2.1 Two weeks after surgery start head-fixing the mouse for 10 min by securing the head bar with pinch clamps for acclimating the mouse to being head fixed.

2.2 Head-fix the mouse three weeks after the surgery for placement of the baseplate.

2.3 Attach a miniscope to a 3D printed holder attached in a micromanipulator (Miniscope Holder, Jove Materials Spreadsheet) attached in a micromanipulator (Figure 3A, B).

**Figure 3:**
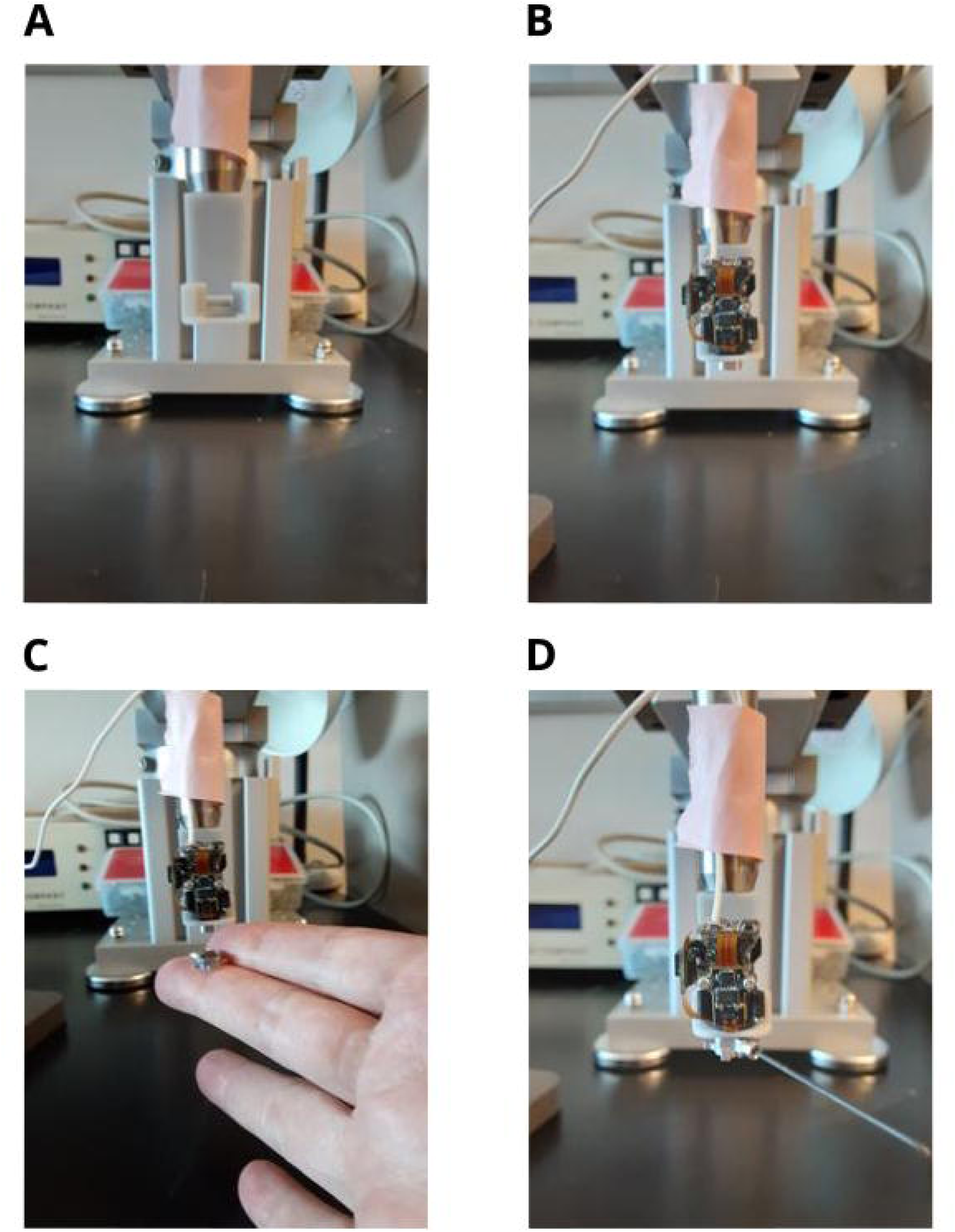
Miniscope base plate placement. A: 3D printed miniscope holder coupled to a micromanipulator. B: Miniscope attached to the holder. C: Attaching the baseplate to the miniscope. D: Tightening the set screw for fixing the baseplate to the miniscope. The set screw is released after cementing the baseplate to the mouse skull.

2.4 Carefully remove the silicone adhesive (Jove Materials Spreadsheet) from the top of the lens using fine tweezers and clean the surface of the GRIN lens using a lens wipe.

2.5 Secure the baseplate attached to the miniscope tightening the set screw on the side of the baseplate (Miniscope baseplate V2, Jove Materials Spreadsheet) (Figure 3 C, D).

2.6 Move end of the miniscope until reaching 100 µm above the GRIN lens.

2.7 Use the sliders to configure the Miniscope settings in the Miniscope software (Miniscope DAQ-QT Software, Jove Materials Spreadsheet) following the steps 2.8 to 2.10.

2.8 Click in the power slider in the software to control the power and keep it around 10%.

2.9 Click in the acquisition rate slider and keep it around 10 Hz.

2.10 The v4 miniscope has an electrowetting lens that can be used for fine focusing. Click in the focus slider to set it at 50% to keep the electro-wetting-lens in the middle of the focal range.

2.11 Use the Miniscope software to view the top of the GRIN lens. It is possible to see the circular shape of the GRIN lens on the screen.

2.12 Bring the miniscope closer to the GRIN lens by adjusting the axial position with the micromanipulator and monitor the image with the computer. Once the top of the GRIN lens is visible, use fine focus up and down on the micromanipulator to reach the best focal plane to visualize cell flashes. Use the pseudo dF/F0 setting in the miniscope software to double check the quality of the cell flashes. The focus is set to allow the maximum number of cells in the field of view with highest attainable fluorescence intensity. At this point blood vessels should be in focus.

2.13 It may happen that a mouse has no transient calcium signals -flashes-. In this case, it is recommended to avoid the next steps (2.15 to 2.17) and verifying the mouse again in the next weeks to double check whether the transient calcium signals occur. A decision can be made to euthanize the mouse if the transient calcium signals never occur two months after the surgery.

2.14 Once the focus is optimized, carefully build a wall of quick adhesive cement between the skull and the bottom of the baseplate. Take care not to cement the miniscope on the baseplate and be very careful not to get any cement on the top of the GRIN lens.

2.15 Once the cement is dry, remove the set screw and carefully remove the miniscope from the baseplate with the micromanipulator.

2.16 Use the base plate cover to protect the GRIN lens.

3. Construction of the Odor Arena

This method delineates an automated odor arena based on the design of Connor et al.^13^ and Gumaste et al.^14^. The complete assembly can be found <u>here</u> (<u>Odor Arena Hardware</u>, Jove Materials Spreadsheet) (Supplementary File 3).

3.1 The chamber dimensions are 50 cm (L) x 50 cm (W) x 25 cm (H) with 2 acrylic walls, an acrylic ceiling, a white expanded PVC floor, and 2 unique walls at the front and rear which facilitate air flow (Figure 4A).

**Figure 4.**
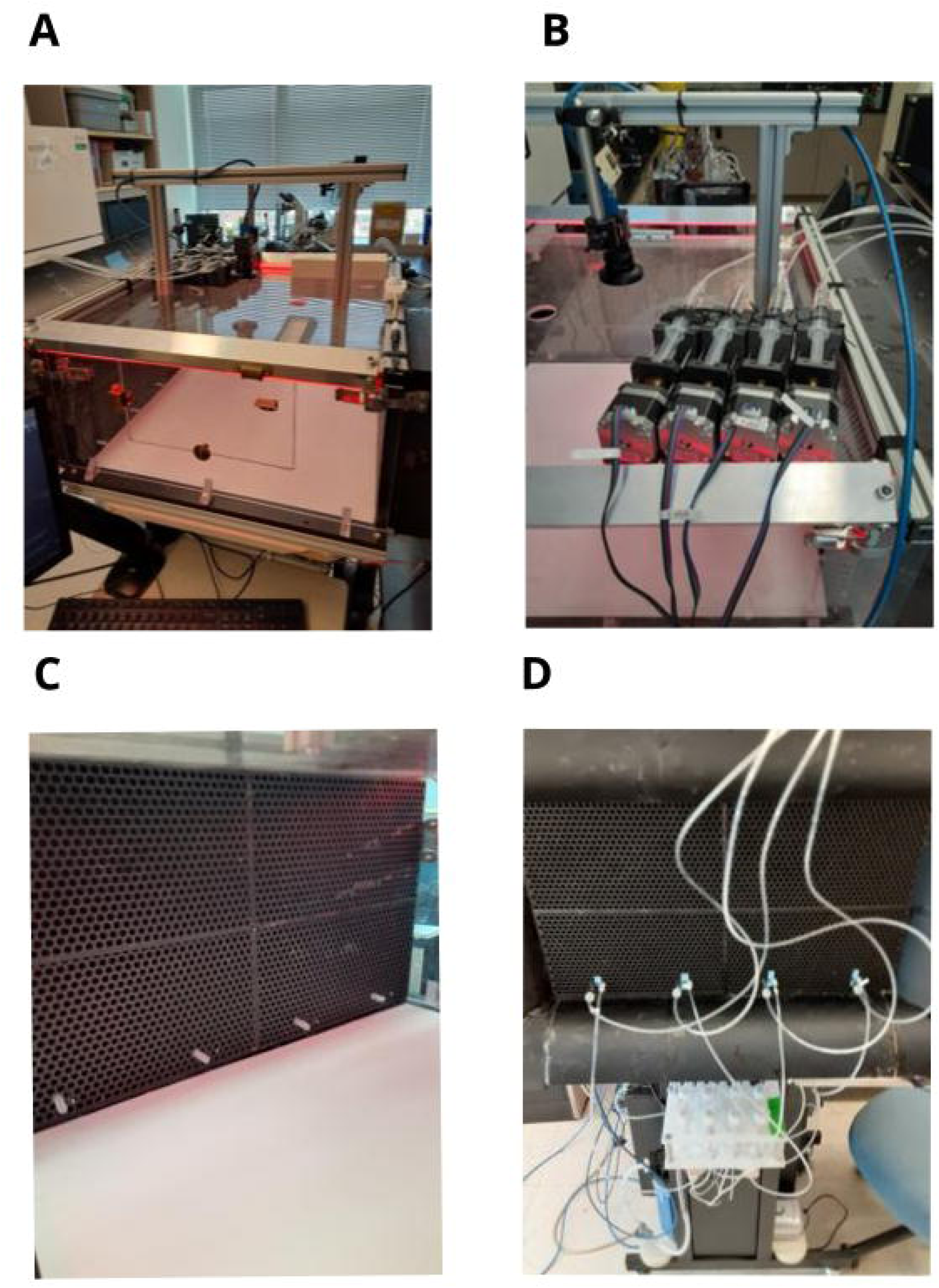
Construction of the Odor Arena. A: Odor arena transparent Chamber. There is a top digital camera for recording the mouse behavior. B: Step motors coupled to syringes control the water delivery to reward the mouse. C: View from the inside of the odor arena showing a honeycomb structure is used to yield a laminar flow and four odor delivery lines. D: Odor delivery system including tubes, valves and odor bottles viewed from the outside of the odor arena.

**Figure 5.**
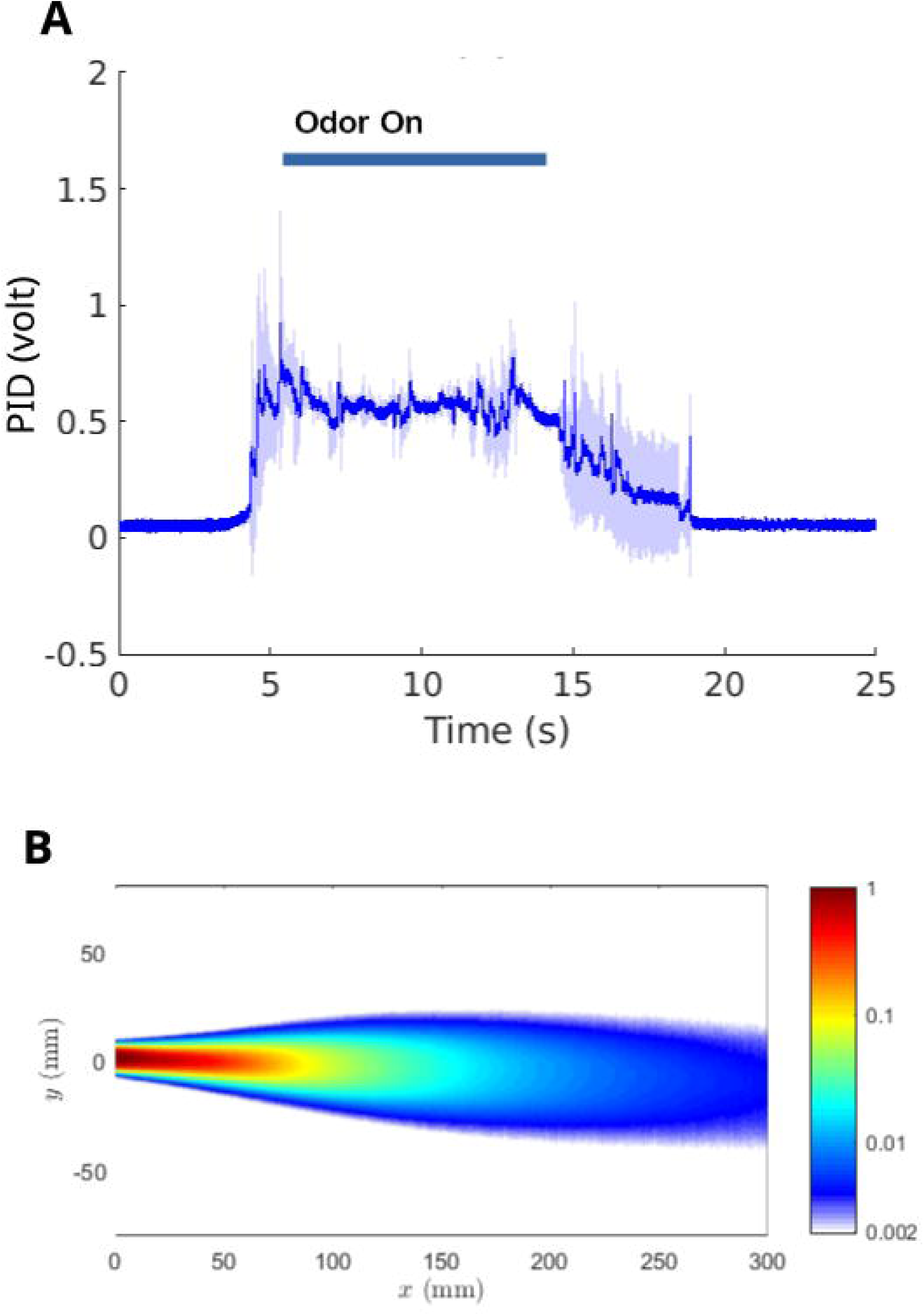
Odor plume recording. A: Odor plume recorded with the photoionization detector (PID). Mean(bold blue)±Standard Deviation (light blue) of five PID traces of an odor plume propagating at 4.23 cm/s in the odor arena. Odorant source at 2cm from the odor arena floor. The sensor head of the PID provides a voltage signal for the gas concentration of the odor plume. B: Laser recording of the odor plume with unbounded flow at 20 cm/s.

3.2 The air suction end at the rear of the arena has a tapered design with a computer fan attached to draw air out of the chamber. A physical knob sets the fan PWM to regulate the rate of air flow. A 3D printed honeycomb structure comprises walls for the front and rear of the odor arena to facilitate laminar flow of air while retaining a mouse inside ^10^. The honeycomb wall at the front of the arena has an external flare for receiving large volumes of air in a non-disruptive manner.

3.3 The water reward is delivered via simple contraptions having a Nema 17 stepper motor coupled to a syringe. An A4988 microstepping driver allows precise control of the dispensed volume (https://github.com/dougollerenshaw/syringe_pump, Figure 4B).

3.4 The air intake end has 4 sets of odor sources paired with water-delivery spouts, with each set positioned 10 cm apart along the x axis to define the ‘lanes’ along which an animal will navigate for a water reward (Figure 4C, D).

3.5 Odor delivery is managed by a system of solenoid valves connected to tubes and odorant bottles (Figure 4D). The valves are powered using a relay board in an arrangement which guarantees that either clean air or odorized air is flowing at all times, and that either one single lane or zero lanes can receive odorized air at any given time. A set of five 12-volt solenoid valves are managed using a relay board containing 4 relays. When all 4 relays are set to ‘off’, the clean-air valve is open by default. When any single odor valve is opened, the clean-air valve is automatically closed. The relay board state is managed using digital outputs from the primary arduino controller. A consumer-grade aquarium air pump supplies air which is restricted to 20 mL/min using a manual air-flow regulator. Using a series of 1/16^th^ inch inner-diameter tubes and splitters, the supply line delivers clean air each of the solenoid valves. Check-valves before and after the odorant guarantee the direction of air flow. The clean air line merges with the odorant lines prior to entering the arena to guarantee that odorized air is purged when the odor lines are closed.

3.6 Animal behavior is monitored at 60 Hz using a single fast digital camera mounted above the arena (Jove Materials Spreadsheet). A 3.5mm fixed focal length C-Series lens was mounted via a C/CS mount adapter, capable of capturing the entire arena (<u>Focal Lens</u>, Jove Materials Spreadsheet).

3.7 The odor arena hardware is managed using custom software developed in Python. A PC connected to a Teensy 4.0 development board provides the means for computer-mediated odor and water delivery. The software integrates the camera and all hardware necessary for setting up experimental parameters and acquiring experimental data (<u>Odor Arena Software</u>, Jove Materials Spreadsheet).

3.8 The digital camera exports a clock signal when recording video frames. The signal is used for post-hoc synchronization with the Miniscope using an USB interface board (Jove Materials Spreadsheet) which records the sync out signals from both systems. During an experiment, the custom acquisition software also creates an events file which contains the important experimental events and the camera frame on which the event occurred. A timestamps file is also created to identify any dropped frames, a rare event.

4. Air speed of the plume measured with a photo-ionization detector

This method detects the time course of the odor plume through a photo-ionization detector (PID) that exposes the gaseous odorant to a high-intensity ultraviolet light that ionizes the odorant molecules. The device’s output detects odorant molecules in the odor plume. This technique allows the estimation of the air speed in the odor arena by comparing the delay to detect the presence of odorants traveling through two locations using the PID.

4.1 Place a fast response miniature PID at two different distances. One location is close and another location far -10 cm apart-from the odor source

4.2 Change the switch of GAIN in the front panel of the PID controller to the position x5.

4.3 Change the switch of PUMP in the front panel of the PID controller to position high.

4.4 Check the LED status light showing the sensor (voltage) output in the front panel of the controller in the absence of odorants.

4.5 Switch the potentiometer OFFSET for zeroing the voltage output in the absence of odorants.

4.6 Turn on the odor valve in the odor arena.

4.7 Measure the delay to detect the odor plume with the PID at each location after opening the valve. This procedure can be performed offline by recording simultaneously the output of the PID and the output of the valve recorded by the odor arena (odor on) with an interface board (Jove Materials Spreadsheet).

4.8 Divide the difference of the delays in each location by the difference of the distance between the two PIDs to calculate the air speed of the plume.

5. Behavioral training mouse in the Odor Arena

This section describes a behavioral task adapted from Findley et al.^4^. The mouse is water restricted the day before to motivate seeking a water reward. The mouse navigates the odor plume (Figure 6B) towards a water spout located at the source of odor release to obtain water reinforcement (3 drops of 10 µL delivered at 1Hz). During the training period the mouse is maintained under the water restriction by having access to up to 2 mL a day. The body weight of the mouse is monitored during the water restriction period and should not be bellow 85% of the original body weight. The mouse receives approximately 1 mL of water per day during the training in the odor arena and is supplemented with an additional 1 mL of water per day in the cage after training. The mouse stays under water restriction for a maximum period of 72 hours. Custom software (Jove Materials Spreadsheet) detects the mouse’s location in real-time (60 Hz) using a simple background-subtraction and blob-localization technique. The user manually sets lane boundaries, the home boundary (the starting location for the mouse at the back of the arena), and the target boundary (near the odor source at the front). Additionally, the user is can decide how the software utilizes these boundaries. For example, the user may deliver odors only when the mouse is behind the home boundary. For the mouse to receive a reward, the user may require it remain within the odorized lane as it navigates to the target (the odor source). Once the mouse crosses the target boundary, it may receive a reward. During training, however, any of these requirements are adjusted by simply editing a ‘yaml’ file designed to be self-explanatory and user-friendly.

**Figure 6.**
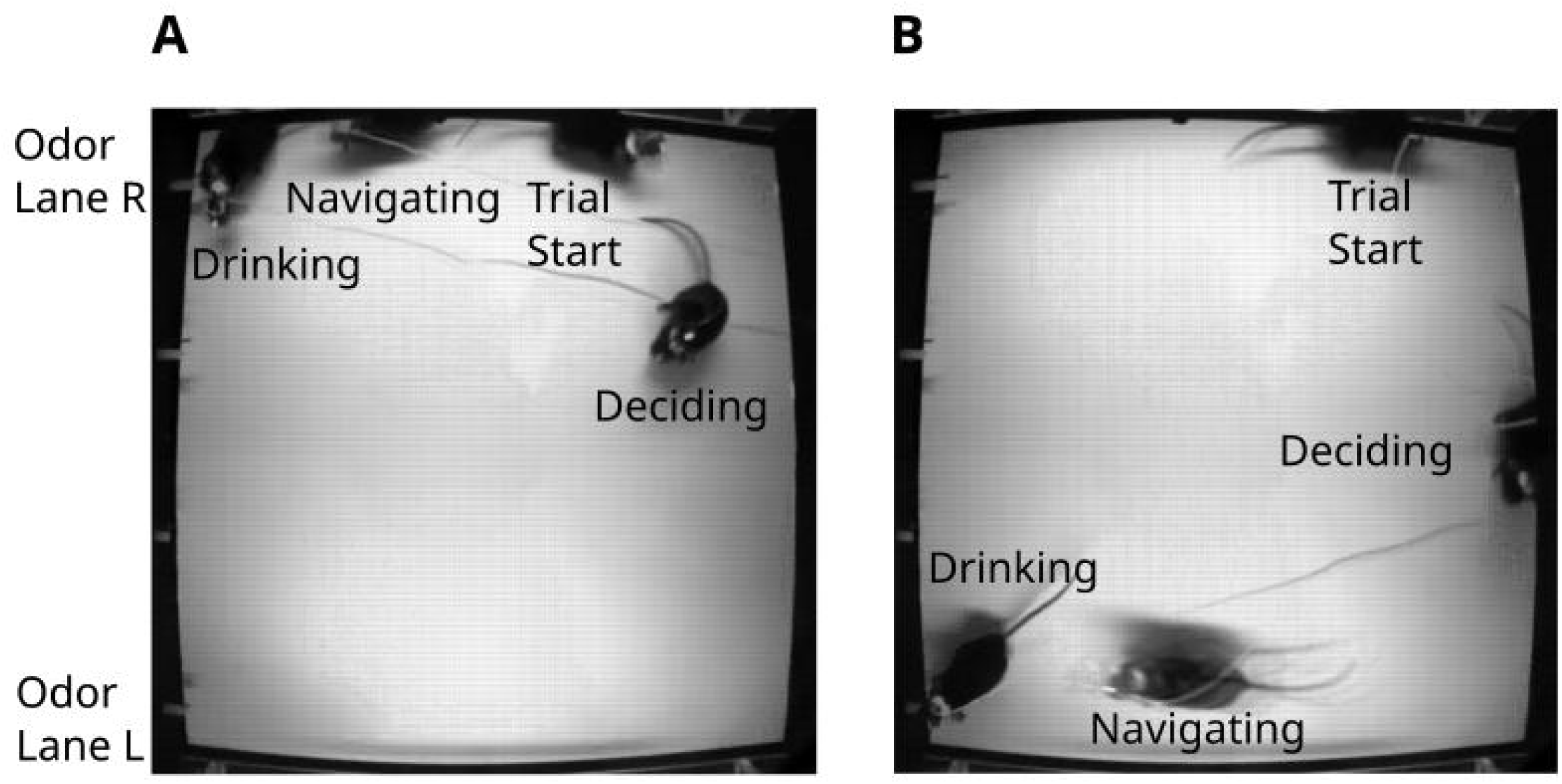
Mouse Behavioral Training. A: Mouse learning to navigate toward an odor plume released on the right lane. The mouse learns to start a trial by going to the back of the odor arena, decide a side to navigate toward the odor plume, and drink a water reward. B: Mouse learning to navigate toward an odor plume released on the left lane. The mouse is rewarded with odor if navigating to the correct lane.

5.1 First, train the mouse to start trials by moving to the back of the arena (defined as the portion of the arena that is 40 cm away from the side where the air flows into the chamber). Wait until the mouse goes to the back of the arena and then manually deliver odor and water in one random lane and let the mouse find the source and drink the water. Repeat the procedure many times to create the association between odor and water (Figure 6A, B). Once the mouse learns to start trials, then use the automated software to deliver odors.

5.2 Train the mouse to do the two-lane odor navigation task using the automated custom software. In this task one of two odor ports is randomly chosen to deliver odor and the mouse is reinforced with water when it arrives to the water spout where odorant is delivered. This protocol used the odorant Isoamyl Acetate diluted at 1% in mineral oil.

5.3 The mouse completes a session of about 20 trials of odor plume navigation in about 40 min. The mouse performs one session per day.

5.4 The trained mouse achieves a percent of correct navigation above random choice in the end of the session (> 65% correct choices). The mouse should achieve the criterion of > 65% correct choices on the two-lane odor navigation task after 3 to 5 sessions of training.

6. Epifluorescence recording of a freely moving mouse in the odor arena

The method describes how to record the neuronal activity of stratum pyramidale (SP) cells in dorsal CA1 by imaging the genetically encoded calcium sensor GCaMP6f expressed in Thy1 mice^9^ by wide-field miniscope imaging during the two-spout odor plume navigation task (Supplementary Movie 1, 2). A typical imaging session takes 40 min allowing the mouse to complete approximately 20 trials of odor navigation. This technique records a mouse for several months.

6.1 Head fix the mouse, place the miniscope on the top of the baseplate using the micromanipulator and tighten the set screw.

6.2 Adjust the electrowetting lens to find the optimal focal plane with the largest number of cells with highest fluorescence intensity.

6.3 Adjust the miniscope power to obtain the optimal dynamic range with a high signal to noise ratio without saturation. This is usually attained imaging dorsal CA1 in Thy1-GCaMP6f mice with miniscope power set around 30% at an acquisition rate of 30Hz.

6.4 Release the mouse inside of the odor arena with the miniscope attached to the baseplate.

6.5 Start acquisition with the interface board to record the TTL output of the digital camera located at the top of the arena and the TTL signal from the miniscope for latter synchronization between behavioral and GCaMP6f video frames. The digital camera records at 60 Hz and the miniscope records at 30 Hz.

6.6 Start recording the miniscope and behavioral movies and turn on the automated software for two spout odor navigation task.

7. Data Preprocessing

This method uses a MATLAB pipeline to process the data. The code is available on GitHub (Synchronization Software, Jove Materials Spreadsheet). NoRMCorre^31^ was used for motion correction, and EXTRACT^32^ to find the ROIs with time-varying fluorescence signals reported as changes in fluorescence normalized by fluorescence between calcium transients (dF/F0).

7.1 Synchronize the odor arena metadata, the digital camera frames and the miniscope frames using the sync signals recorded by the interface board by running the MATLAB code Synchronize_Files_JOVE.m available on GitHub (Synchronization Software, Jove Materials Spreadsheet).

7.2 Perform motion correction of the synchronized miniscope frames using NoRMCorre (Motion Correction Software, Jove Materials Spreadsheet).

7.3 Find the ROIs with time-varying dF/F0 signals using EXTRACT (ROI Extraction Software, Jove Materials Spreadsheet).

7.4 Separate the data into trials (Synchronization Software, Jove Materials Spreadsheet).

7.5 Label each trial either as a hit or a miss based respectively on the mouse’s correct or wrong navigation behavior.

7.6 Use Behavior Ensemble and Neural Trajectory Observatory (BENTO)^33^ to visualize the behavior and ROIs of each separate trial (Figure 7 A, B, Supplementary Movie 3).

**Figure 7.**
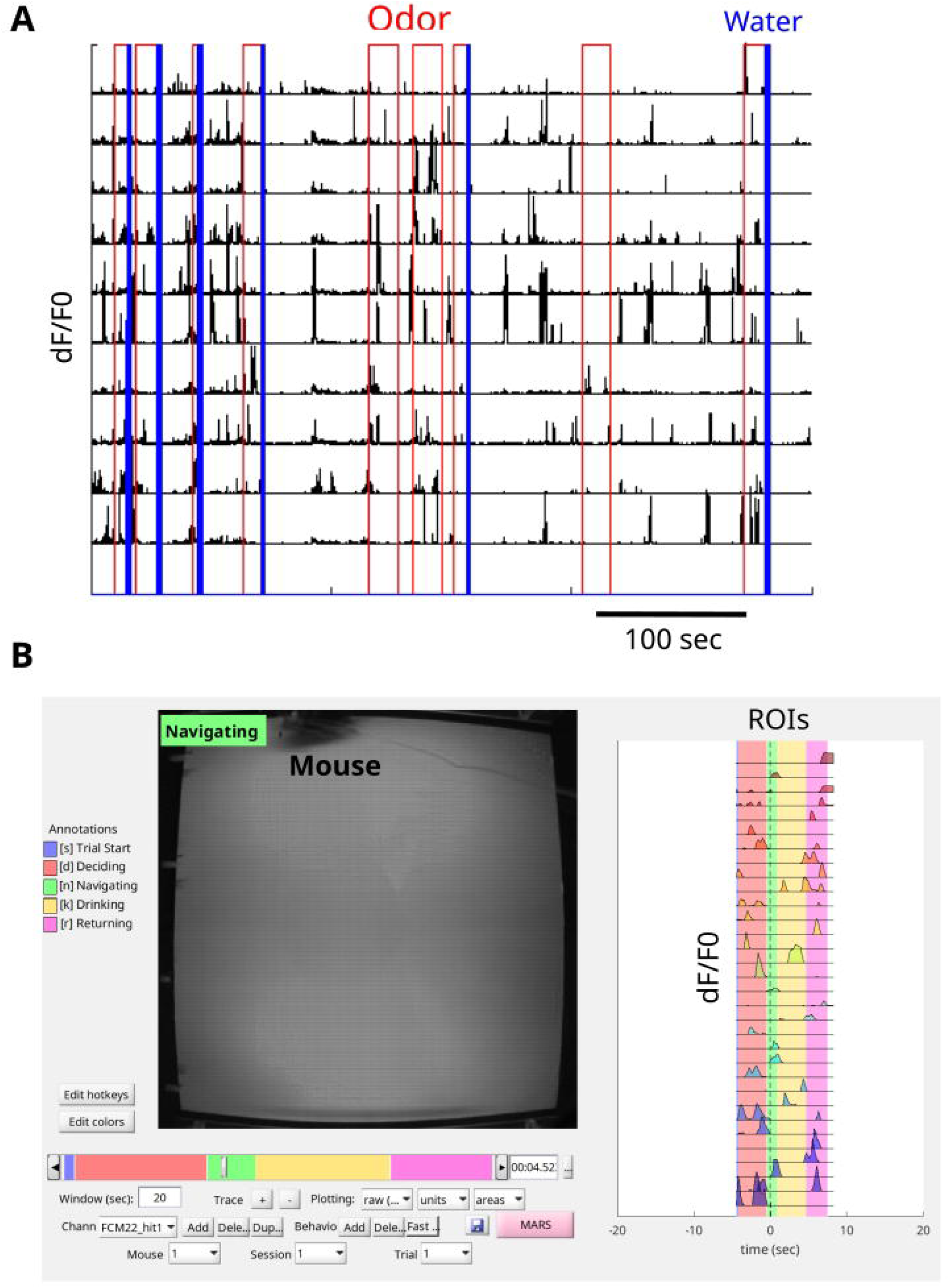
Pre-Processing the Data A: Calcium traces synchronized with the odor arena events for many trials. Each single trial starts with the odor delivery in red and the rewarded trials end with a water delivery pulse in blue. The dF/F0 (unitless) calcium traces for each ROI are shown in black. Each line indicates a ROI. An interface board is used for recording the TTL outputs of the top camera on the odor arena and the miniscope camera for synchronizing the frames. Normcorr is used for correcting the movement noise from the miniscope frames and Extract is used to find the ROIs and extract the dF/F0 calcium traces. B: Representative single trial of a mouse navigating a plume. The simultaneous visualization of the synchronized behavior (left panel) and dF/F0 calcium traces (right panel) from each ROI of a single trial are observed with BENTO.

8. Data Analysis – Decoding Spatial Position from Brain Signals

This method uses machine learning to decode the mouse’s X and Y positions in the arena from the dCA1 ROIs^7^. The MATLAB code is available (Decoding Brain Signals Software, Jove Materials Spreadsheet) in https://github.com/restrepd/drgMiniscope.

8.1 The software drgDecodeOdorArenav2.m takes as input the EXTRACT.mat output file and the odor arena metadata file. The GitHub repository provides these two files for an example (Synchronization Software, Jove Materials Spreadsheet):

dFF_file=’20220804_FCM22_withodor_miniscope_sync_L1andL4_ncorre_ext.mat’; arena_file=’20220804_FCM22withodor_odorarena_L1andL4_sync.mat’;

8.2 drgDecodeOdorArenav2.m creates a dataset divided by within trial data containing the ROI signals and metadata (X and Y positions of the mouse, odor spout location, water delivery, etc) for each trial. The code also analyzes decoding for data between trials.

8.3 The code uses fitrnet to train an artificial neural network with the dF/F0 data for all ROIs for all trials but one to predict x and y positions and predict the position of the trial left out using a leave one out procedure.

8.4 The neural network returns the X and Y positions as output. Use the trained neural network to make predictions. Input left out trials not used to train the network to make predictions of the X and Y positions of the mouse in the arena from the ROIs (Figure 8A, B).

**Figure 8.**
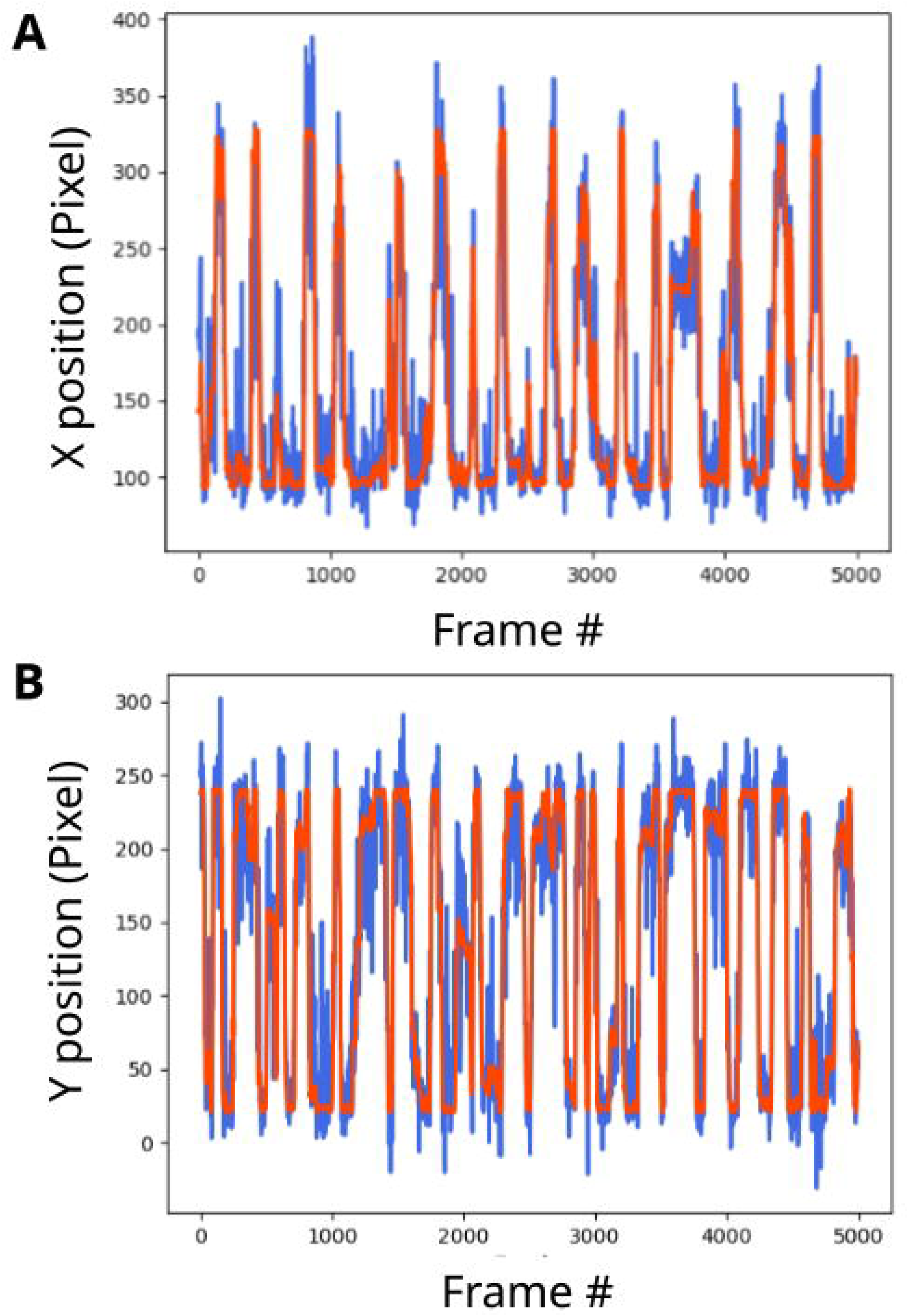
Decoding the position of the mouse from the CA1 signals. Decoding the X and Y position of the mouse from the CA1 ROIs. The decoding prediction is shown in blue and the mouse ground true position is shown in red. The predicted traces are strongly correlated with the ground truth. A: Decoding the X positions from the ROIs (Pearson Correlation Coefficient = 0.88). B: Decoding the Y positions from the ROIs (Pearson Correlation Coefficient = 0.88).

## REPRESENTATIVE RESULTS

Using this procedure, allows visualizing and recording dCA1 GCaMP6f fluorescence transients in mice navigating the odor arena to find the source of odorants (Figure 6 A,B, Supplementary Movie 1, 2). The fluorescence images are motion-corrected with NoRMCorre, and EXTRACT is used to extract the ROIs. In addition, recording with an interface board allows to synchronize the dF/F0 signals from the ROIs with odor and water delivery events in the odor arena (Figure 7A), as well as with movement of the mouse in the odor arena (Figure 7B, Supplementary Movie 3). The representative result of the mouse navigating the odor plume includes a large number of calcium transients during the task (Figure 7 A, B). In addition, it is possible to inspect how the calcium responses are aligned with the presence of odor and water reward (Figure 7A). The visualization of single trials with BENTO provides information about the calcium responses at different stages of the trial, including trial start, deciding, navigating, drinking, and returning to the back of the arena (Figure 7B). The method communicated valuable insights regarding the link between CA1 calcium responses and mice behavior during an odor-oriented navigation task.

The PID recordings can provide crucial information about the odor plume and the air speed of the plume. The representative result shows an increase in the PID response after opening the valve to release the odor plume inside of the odor arena (Figure 5A). Furthermore, the protocol yields decoding of the X and Y positions of the mouse from the dF/F0 signals of the dCA1 ROIs (Figure 8). This technique predicts the spatial location of the mouse during the odor-plume navigation task based on the CA1 responses, which is relevant to better understanding how the CA1 neurons process odor and spatial information. Decoding the trajectory of the mouse from neuronal ensemble calcium signals in dCA1 is significant because it reveals how neurons in dorsal CA1 represent a cognitive map of odorant and spatial information to perform the complex task of odor plume navigation. The method has been expanded to different ROIs that behave exclusively as place cells and other cells that respond to the odor stimulus. Successful decoding of the trajectory of the mouse from neuronal ensemble signals can be confirmed by the strong correlation between the decoding prediction and the ground true X and Y positions of the mouse.

## SUPPLEMENTARY FILES

Supplementary Movie 1:

Representative example of synchronized behavioral and miniscope frames of a mouse navigating toward an odor plume in the left lane. A: Behavioral frames of a mouse wearing a miniscope and navigating inside of the odor arena. B: Miniscope frames of the mouse showing the raw calcium transients recorded through the GRIN lens.

Supplementary Movie 2:

Representative example of synchronized behavioral and miniscope frames of a mouse navigating toward an odor plume in the right lane. A: Behavioral frames of a mouse wearing a miniscope and navigating inside of the odor arena. B: Miniscope frames of the mouse showing the raw calcium transients recorded through the GRIN lens.

Supplementary Movie 3:

BENTO display of the processed data of mouse behavior and brain signals. Left panel: the mouse is navigating toward the right lane in the arena. The behavior annotations are showed in different colors. B: dF/F0 calcium signals of the mouse navigating.

Supplementary File 1:

Files for the 3D printed nose cone to perform isoflurane anesthesia.

Supplementary File 2:

Files for the head bar for head-fixing the mouse.

Supplementary File 3:

Detailed layout of the odor arena.

## DISCUSSION

This protocol meticulously outlines the steps to record place-cells and odor-responsive cells in the dCA1 area of the hippocampus of mice navigating an odor plume. The critical steps in the protocol include stereotaxic surgery, placement of the miniscope baseplate, construction of the odor area, checking the plume in the odor arena, behavioral training, miniscope recording of the freely moving mouse, data pre-processing, and data analysis. Additionally, the protocol explains the process of decoding the mouse trajectory from the dCA1 recordings.

A critical step in stereotaxic surgery is to follow the coordinates relative to Bregma to place the GRIN lens at the correct location. A limitation of the method is the delay between the surgery and the time to start observing GCaMP6f signals, which can take two to four weeks. The mouse should be ready for use after this critical period. A difference between the current protocol and previous protocols ^24, 24–26, 26, 27^ is using Thy1-GCaMP6f mice already expressing GCaMP6f in CA1 instead of injecting AAV-GCaMP6f virus in the hippocampus. It saves time during the surgery and does not require a wait time for the expression of the AAV virus in the brain. In addition, this protocol does not aspirate the brain and uses isoflurane instead of ketamine/xylazine, which provides a better control of the dose to prevent overdoses. A limitation of the method is the optical aberration of the GRIN lens, which limits the field of view. A crucial step for the baseplate placement is to keep the electrowetting lens at the center to prevent a limitation in adjusting the Z plate after cementing the baseplate on the head. A limitation of the construction of the odor arena is the system’s complexity, which can take several months and may require the assistance of an engineer. A limitation of the behavioral training is that mice may prefer one side of the odor arena against another. One way to overcome this problem is to alternate trials rewarding the mouse for navigating toward the left and right lanes.

A critical step for checking the plume is to keep the PID needle aligned with the odor source to detect the plume’s path. A crucial step in miniscope recording a freely moving mouse is to ensure that the mini-coax wire does not tangle during the task, which can be prevented with a commutator. Helium balloons can be used to prevent the mini-coax wire from coming in front of the mouse. It is critical for the data pre-processing to synchronize the TTL pulses of the digital top camera of the odor arena and the miniscope camera. For the EXTRACT procedure, it is recommended to use non-negative processing to extract the ROIs and dF/F0 signals better. EXTRACT yields a visual inspection of the traces for each ROI to exclude bad ones. It is critical to decode the X and Y positions of the mouse from the ROIs to have a large dataset with hundreds of epochs for better training the artificial neural network.

The significance of this freely moving recording method concerning existing head-fixed methods is studying the mouse behavior in an ethologically relevant context with the proper head movement for navigating a complex odor plume. This method is applicable to investigate the dynamic role of dCA1 neurons in complex odorant navigation. Furthermore, the procedure is not restricted to hippocampus or olfaction. Other potential applications of the technique include studying the role of different brain areas and sensory modalities, including possible applications in visual navigation by using LEDs to indicate the rewarded lanes. In addition, this method can be potentially applied in real-time closed-loop experiments in which decoding neural calcium from population triggers neurostimulation or sensory feedback ^20, 34–36^.

## Supporting information

Jove Materials

## ACKNOWLEDGMENTS

This research was supported by the US National Institutes of Health (NIH UF1 NS116241 and NIH R01 DC000566), and the National Science Foundation (NSF BCS-1926676). The authors thank Andrew Scallon for helping setting up the Odor Arena chamber.

