## Supplementary material for "Miniscope Recording Calcium Signals at Hippocampus of Mice Navigating an Odor Plume": Jove Materials

| Name of Material/ Equipment | Company | Catalog Number | Comments/Description |
|---|---|---|---|
| Arduino Micro | Arduino | Micro | |
| Biocompatible Methacrylate Resin | Parkell | S380 | C&B-Metabond Adhesive Luting Cement |
| Data Acquisition System (DAQ) | Labmaker | NA | DAQ for UCLA Miniscope V4 |
| Decoding Brain Signals Software | CU Anschutz | | https://github.com/restrepd/drgMiniscope |
| Dental Drill | Osada | LHP-6 | AZ210015 |
| Dental Drill Box | Osada | XL-230 | 30000 rotations per minute |
| Digital stereotaxic instrument | Stoelting | 51730D | Mouse Stereotaxic Instument, #51904 Digital Manipulator Arm |
| Drill Bit | FST Fine Science Tools | 19007-05 | Tip diameter 0.5mm |
| Fast Digital Camera | Edmund Optics | BFS-U3-63S4C | FLIR Blackfly S |
| Focal Lens | Edmund Optics | C-Series | 3.5 mm |
| GRIN lens | Inscopix | 1050-004595 | 1mm diameter and 4mm length |
| GRIN lens Holder | UCLA | | http://miniscope.org/index.php/Surgery_Protocol |
| Liquid Tissue Adhesive | 3M | 1469C | Vetbond Tissue Adhesive |
| Low-Flow Anesthesia System for Mice | Kent Scientific Corporation | SomnoSuite | https://www.kentscientific.com/products/somnosuite/ |
| Low Toxicity Silicone Adhesive | WPI – World Precision Instruments | Kwik-sil | |
| miniPID Controller | ASI – Aurora Scientific Inc. | Model 200B | Fast-Response Miniature Photo-Ionization Detector |
| Miniscope V4 Holder | UCLA | NA | https://github.com/Aharoni-Lab/Miniscope-v4/tree/master/Miniscope-v4-Holder |
| Miniscope V4 | Labmaker | NA | https://www.labmaker.org/products/miniscope-v4 |
| Miniscope Base Plate V2 | Labmaker | NA | https://www.labmaker.org/products/miniscope-v4-base-plates |
| Miniscope DAQ-QT software | UCLA | | https://github.com/Aharoni-Lab/Miniscope-DAQ-QT-Software/ |
| Motion Correction Software | CU Anschutz | | https://github.com/restrepd/drgMiniscope |
| Odor Arena Hardware | Custom Made | 3D Model | https://www.dropbox.com/scl/fo/lwtpqysnpzis32mhrx3cd/ADomsxy |
| Odor Arena Software | CUAnschutz | | https://github.com/wryanw/odorarena |
| Odorant Isoamyl Acetate | Aldrich Chemical Co | 06422AX | Diluted at 1% in odorless mineral oil |

| | | | |
|---|---|---|---|
| RHD USB Interface Board | Intan Technologies | C3100 | Product discontinued. Alternatively use another equivalent boa |
| ROI Extraction Software | CU Anschutz | | https://github.com/restrepd/drgMiniscope |
| Sutter Micromanipulator | Sutter Instrument Company | MP-285 | |
| Synchronization Software | CU Anschutz | | https://github.com/fsimoesdesouza/Synchronization |
| Thy1-GCaMP6f mice | Jackson Laboratory | IMSR_JAX 028281 | C57BL/6J-Tg(Thy1-GCaMP6f)GP5.12Dkim/J) |

, 3-Axes, Add-On, LEFT

-variant-2-pack-of-10
wiki

hxu42sqDmTBl2O6k?rlkey=b3l4809eradundt5l3iz0gq74&dl=0

ırd.
