## Supplementary figures and images for "Miniscope Recording Calcium Signals at Hippocampus of Mice Navigating an Odor Plume"

### Supplementary Fig3

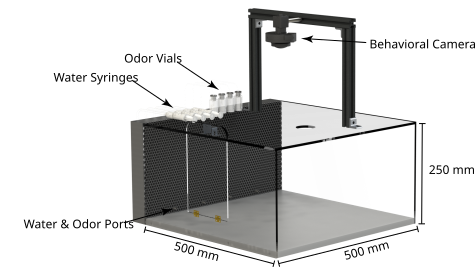
